# Phylogenetic analysis of pigeon paramyxovirus type 1 (PPMV-1) detected in the British Isles between 1983 – 2023

**DOI:** 10.1101/2025.05.09.652871

**Authors:** Alexander M. P. Byrne, Benjamin C. Mollett, Ian H. Brown, Joe James, Ashley C. Banyard, Craig S. Ross

## Abstract

Newcastle Disease (ND), caused by virulent strains of avian paramyxovirus type-1 (APMV-1), is one of the most important poultry diseases globally due to its economic impact and endemicity in lower- and middle-income countries. A variant of APMV-1 is endemic in Columbiformes (pigeons and doves) worldwide and is commonly termed pigeon paramyxovirus-1 (PPMV-1). Since its initial detection in the 1980s, PPMV-1 has caused numerous ND outbreaks in poultry, including in high income countries, and was the causative agent for the last ND outbreak in the British Isles in 2006. Here, we have undertaken sequencing of PPMV-1 isolates between 1983 and 2023 and define three distinct genotypes of PPMV-1 being present in the British Isles. Analysis of the contemporary VI.2.1.1.2.2 genotype, demonstrated likely incursion from mainland Europe, whilst this genotype has subsequently spread across China, with detection also occurring in Australia. The presence of a virulent fusion-gene cleavage site in sequences highlights the continued risk to poultry from PPMV-1 genotypes which were detected in pigeons and doves across the British Isles.

## Introduction

Newcastle Disease (ND) is caused by infection of poultry with virulent strains of avian paramyxovirus-1 (APMV-1) commonly termed Newcastle Disease Virus (NDV). NDV is one of the most important poultry pathogens due to the potential high mortality and subsequent economic losses. Outbreaks of ND are subject to official controls as prescribed by the World Organisation for Animal Health (WOAH) (World Organisation for Animal Health (WOAH) 2021) and both national and international legislation. Alongside the economic losses from culling activities to manage the disease following outbreaks, international trade embargos may be placed upon countries where ND is detected, which results in further significant economic losses.

Virulent APMV-1 is endemic in Columbiformes (pigeons and doves) globally and therefore poses a constant threat to poultry and wild bird species. *Columbiformes* exist as a single family called *Columbidae*, consisting of 41 groups of pigeons and doves, and representing over 300 species. *Columbiformes* have a wide geographical distribution and are regularly seen in every continent except Antarctica. All domestic pigeons represent a single species (*Columba livia domestica*) which were domesticated from wild common pigeons (also referred to as wild rock doves) (*Columba livia*). While small populations of truly wild common pigeons still exist in a few areas, the vast majority of common pigeons seen in the wild have established from domestic escapees (also referred to as feral pigeons or city pigeons), which have high connectivity with human settlements, including poultry farms (Gibbs 2010; Graziosi et al. 2024), whilst racing and show pigeons are in constant contact with humans and are a present in human settlements, and therefore may be in contact with poultry in both domestic and commercial settings. APMV-1 was first isolated in Columbiformes in the 1970s (Kaleta, Alexander, and Russell 1985), whilst the first case in Europe was detected in Italy 1982 (Biancifiori and Fioroni 1983), before rapidly spreading across Europe and being detected for the first time in the British isles in early 1983 (D J Alexander, Russell, and Collins 1984). These Columbiform origin APMV-1s are both antigenically and genetically distinct from commonly derived poultry APMV-1, and due to this distinction, have been named pigeon paramyxovirus type-1 (PPMV-1). Detection of PPMV-1 within the European Union (EU) and British Isles has continued (Dennis J Alexander 2011) with 287 detections in Columbiformes in the EU and other European countries in 2019 alone (AI-ND EURL 2019). Although PPMV-1 is endemic in Columbiformes world-wide, the feral nature of pigeon and dove populations means that prevalence is difficult to determine. Analysis in Switzerland suggested that between 5 -17% of common pigeons are actively infected with PPMV-1, depending on season, with little difference observed due to age or sex (Annaheim et al. 2022), whilst analysis in China suggested an infection rate of 0.5 – 3.19% in live bird markets (Yu et al. 2022).

Phylogenetically, APMV-1 is split into two classes. Class I APMV-1s are regularly detected in wild waterfowl with occasional spill over into poultry (Dimitrov et al. 2016). In contrast, Class II APMV-1s include the vast majority of NDV’s and are predominantly detected in domestic poultry, although isolates are also detected in a wide range of wild birds (Dimitrov et al. 2016). A critical determinant of virulence is the fusion gene (F), and genetic analysis of F is used to genotype APMV-1 due to its heterogeneity and therefore, the best target for investigating genetic relationships. There are currently 20 distinct genotypes based on full F-gene sequence (Dimitrov et al. 2019), with genotypes VI, XX and XXI being associated with PPMV-1 isolates (Dimitrov et al. 2019). Previous analysis using only partial F-gene sequence, a 374 bp fragment, classified PPMV-1s as lineage 4 (Aldous et al. 2003) which were subsequently designated genotype VI when the whole 1,662 bp of F-gene was included (Diel et al. 2012). The presence of a virulent F-protein cleavage site is one of the WOAH determinants of NDV when detected in poultry, along with an intracerebral pathogenicity index (ICPI) score of > 0.7 (World Organisation for Animal Health (WOAH) 2021). The presence of the virulent F-gene cleavage site commonly observed in PPMV-1 isolates would be defined as NDV if they were detected in poultry.

Following the isolation of PPMV-1 from racing pigeons in England in 1983, significant numbers of cases were detected annually (D. J. Alexander et al. 1997). To attempt to reduce the impact of PPMV-1 on racing pigeons, the Racing Pigeons (Vaccination) Order 1994 was introduced in the United Kingdom (Department for Environment Food and Rural Affairs 2019). This designated that all birds attending events, such as pigeon shows or races, must be vaccinated against PPMV-1. Measures to control PPMV-1 infection in racing and show pigeons differ to normal measures of humane killing of infected poultry, even though they are considered captive birds, as they are routinely vaccinated, unlike wild Columbiformes. In Great Britain (GB), these control measures involve the isolation of infected birds for the duration of clinical signs plus 60 days following the resolution of clinical disease before racing or movement can resume (Department for Environment Food and Rural Affairs 2019). However, the threat of spread of PPMV-1 from Columbiformes (both feral and domestic) into poultry settings remains. This has been previously observed within the UK, with the first UK poultry outbreak derived from PPMV-1 likely due to contamination of poultry feed caused by a pigeon infestation which resulted in 23 separate outbreaks across GB, mainly in laying hens in 1983 (D. J. Alexander, Parsons, and Marshall 1984). Further outbreaks linked to PPMV-1 in GB were detected in 550 breeding pheasants in 1996 where nervous signs and decreased egg production were observed. Viruses isolated from these premises had ICPI scores of 0.91 and 1.19 respectively, confirming ND (D. J. Alexander et al. 1997). The most recent ND outbreak in GB occurred in grey partridges (*Perdix perdix*) in 2006 and was hypothesised to be due to spillover of PPMV-1 from feral pigeons (*Columba livia forma urbana*) roosting in loft spaces above the courtyard in which grey partridges were housed (Irvine et al. 2009). This virus was determined to be lineage 4b APMV-1, the same virus seen circulating contemporaneously in GB pigeons; the ICPI score of 1.01 of this virus confirmed ND (Aldous et al. 2014). Although standard practice is to cull all birds on the site, due to the presence of 126 rare breed birds (from 34 species) placed on the International Union for Conservation of Nature (IUCN) Red List of Threatened Species, some birds were spared. Similar rules were implemented as for positive pigeon cases, with a 60-day quarantine and vaccination mandatory. This however highlights the potential risk of PPMV-1 incursion into poultry premises.

Analysis of PPMV-1 detections in GB has been undertaken previously, initial analysis of 178 F gene sequences from viruses detected between 1983 - 2002 determined that the majority of GB isolates were of the 4bi group (VIb, VI.1) (Aldous et al. 2004). Analysis of a further 63 viruses, of which 43 were detected between 2003 - 2010 showed that these more recent viruses were now in the 4bii d-f group (4bii d, Vie/f, VI.1.2.2.1 /VI.1.2.2.2, 4bii e, Via/n, VI.2.1.1.1, 4bii f, VI j/k VI.2.1.1.2.1/ VI.2.1.1.2.2) (Aldous et al. 2014), so distinct changes in the genotype of the UK circulating PPMV-1 had been detected over the eight year period. When eleven PPMV-1 viruses detected in Belgium between 1998 – 2011 were analysed, a further change in PPMV-1 genotype was observed (Meulemans et al. 2002). Analysis of PPMV-1 from China has also revealed differing genotypes over time, with genotypes VI.2.1.1.2.1, VI.2.1.1.2.2. and VI.2.2.2 being detected (Z. Wang et al. 2024; Yu et al. 2022; Zhan et al. 2021), whilst genotype VI.2.1.1.2.2 has been detected in Australia (Shan et al. 2021).

Due to the continuing changes in PPMV-1 genotypes observed world-wide, alongside the likelihood of significant knowledge gaps due to under-sampling of feral populations, and the dynamic nature of those populations, whole genome and full F-gene sequencing of historic and contemporary PPMV-1s from the British Isles was undertaken. These sequences were analysed to re-evaluate the diversity of PPMV-1 circulating in the British Isles with increased resolution, to propose the most likely geographical area from which these originated and to increase the resolution of classification of historical PPMV-1 isolates.

## Materials and Methods

### Identification and virus isolation

Virus isolates were propagated using 9-11 day old embryonated fowls’ eggs originating from a commercial specific pathogen free (SPF) flock under Home Office project license number P5275AD31. Isolates were supplied by the National and International Reference Laboratories for Newcastle Disease at APHA-Weybridge. Forty-nine samples were isolated from Columbiform tissues or swabs and analysed for the presence of PPMV-1 by the national reference laboratory by positive selection using monoclonal antibodies specific to PPMV-1 (D J Alexander, Russell, and Collins 1984), or by RT-PCR (Sutton et al. 2019) and subsequent Sanger sequencing (Collins, Bashiruddin, and Alexander 1993). Also included were a lineage 4 (now genotypes VI) isolate from infected grey partridges (*P. perdix*) isolated during the last UK ND outbreak in 2006 (Irvine et al. 2009) and a poultry isolate from Iraq from 1968 (AG68). The 49 Columbiform-derived isolates analysed in this study are deemed to be representative of the circulating UK PPMV-1 isolates from 1983 – 2023 are described in Table 1

**Table 1:**
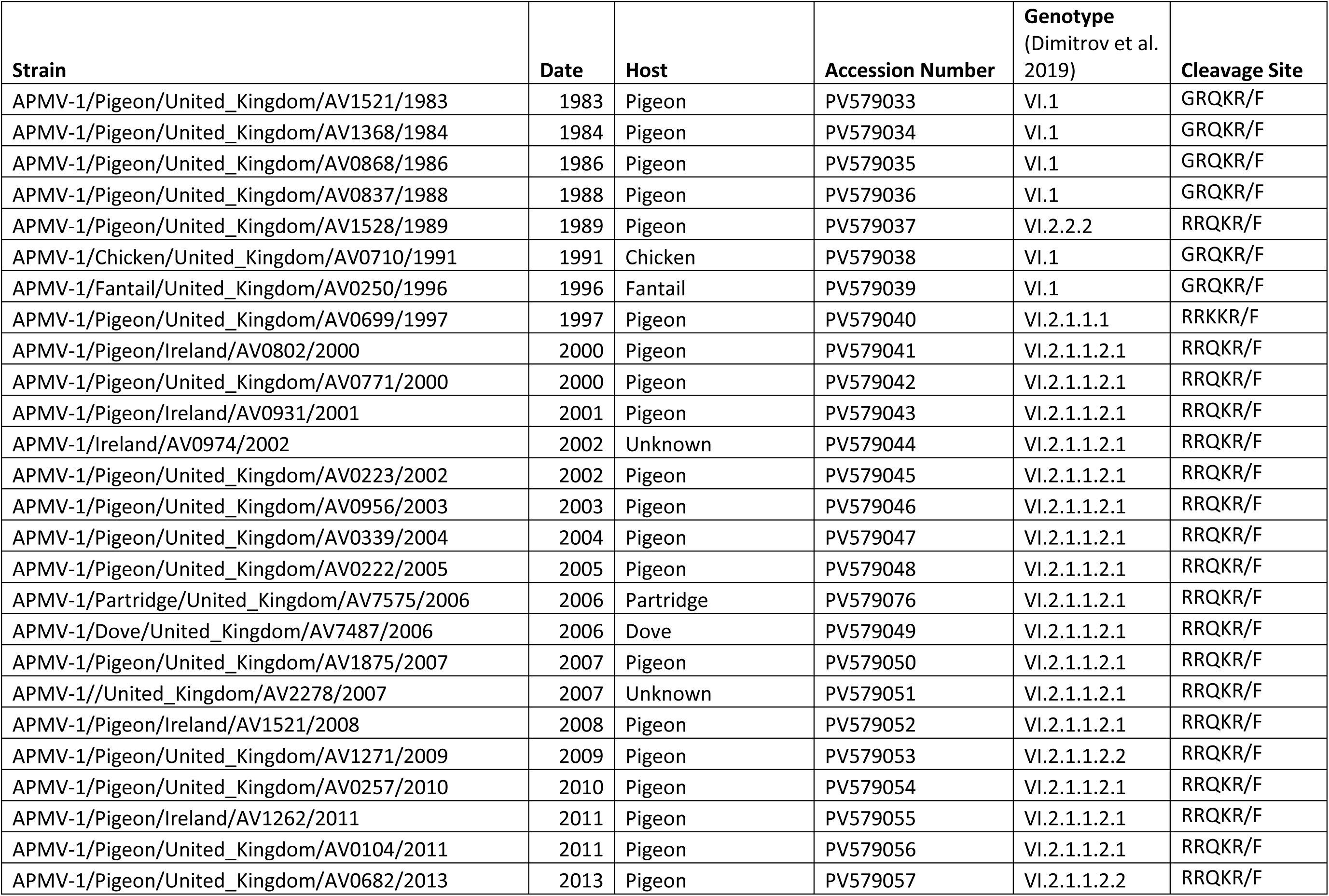
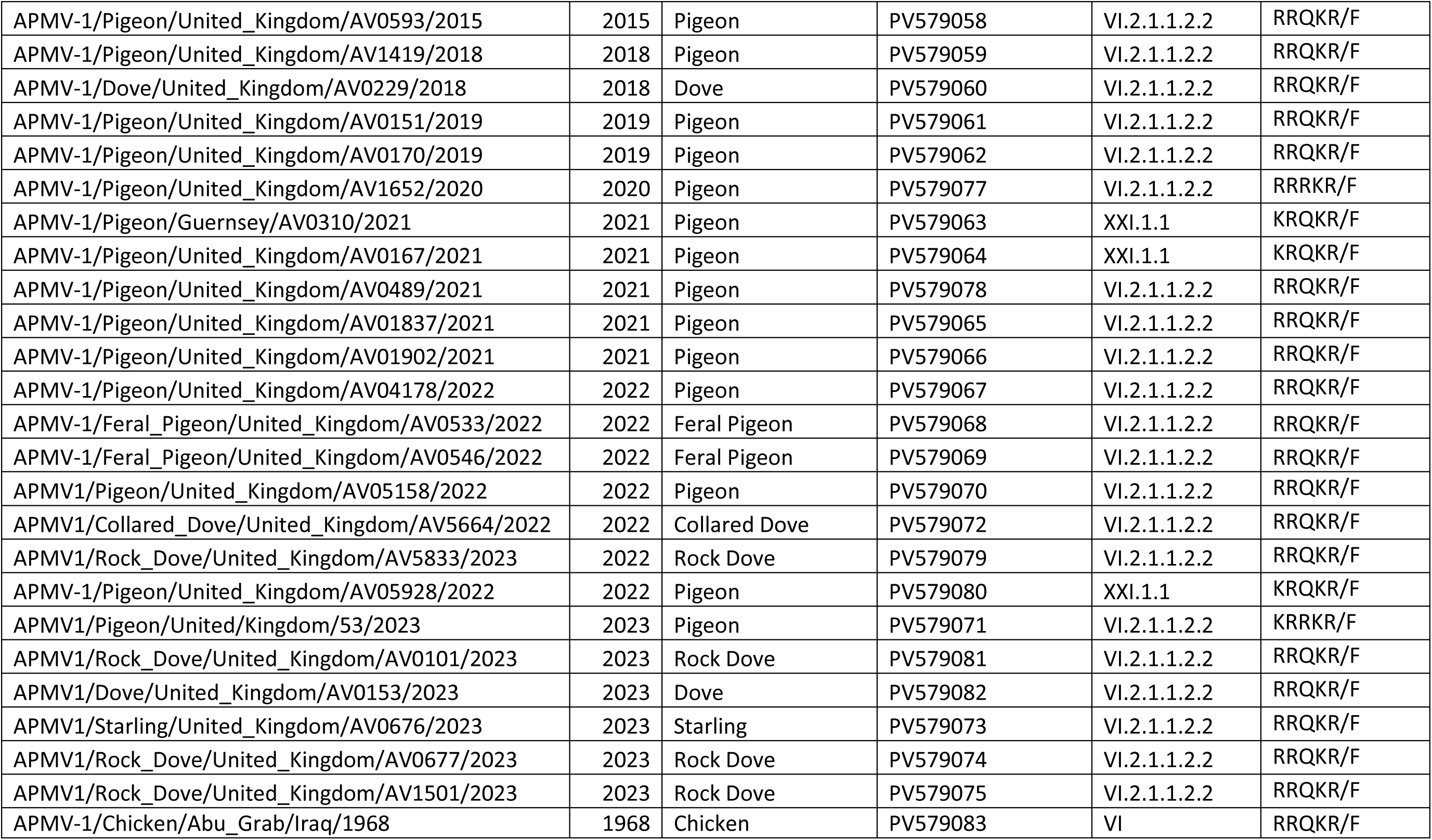
PPMV-1 isolates sequenced and analysed during this study. Isolates, year of isolation, species isolated from, NCBI accession number, predicted APMV-1 genotype and the F-gene cleavage site motif.

### Whole genome sequencing and F-gene analysis by Oxford Nanopore Technologies

Whole genome sequencing (WGS) of selected isolates was carried out as previously described (Reid et al. 2023). Full F-gene fragments were generated for Sanger sequencing and Oxford Nanopore Technologies (ONT) sequencing by amplifying approximately 10 ng viral RNA using OneTaq® One-Step RT-PCR Kit (New England Biolabs) using primers PSMF1 (5’-AGTGACYGCYGAYCAYGAGGT-3’) and POSF1 (5’-TCCGRAAWAYYARGCGCCATGT-3’), with thermocycler conditions of 48°C, 30 minutes; 94°C, 1 minute; followed by 40 cycles of 94°C, 15 seconds; 55°C, 15 seconds; 68°C, 2.5 minutes, followed by 68°C, 5 minutes; and then held at 4°C, generating a PCR product of 2,188 bp incorporating 194 bp of 3’ M-gene, the whole 1792 bp F-gene and 161 bp 5’ of HN, plus intergenic sequences. PCR fragments were purified following gel electrophoresis. Fifty nanograms of PCR product was used for subsequent library preparations. DNA end repair was conducted with NEBNext Ultra II End Repair/dA-Tailing Module (NEB, Hitchin, UK) and purified with Ampure XP beads (Beckman Coulter). The ONT sequencing library was prepared using the Native Barcoding Kit 24 V14 (Oxford Nanopore Technology) and NEB Blunt/TA Ligase Master Mix (NEB). Sample libraries were sequenced with a Minion R10.4.1 flow cell. Sequences were analysed using an in-house pipeline adapted for nanopore sequence data as described previously (Byrne et al. 2023) Coverage of sequences by Illumina, Sanger sequencing or ONT are shown in table S1. Sequences were submitted to NCBI and can be accessed with accession numbers PV579041 - PV579083.

### Evolutionary and geographic analysis

F-gene sequences used in the analysis were downloaded from the ND Sequence consortium (https://github.com/NDVconsortium/NDV_Sequence_Datasets) and from NCBI or BV-BRC databases. Initial phylogenetic analysis was carried out as described previously (Reid et al. 2023).

Bayesian phylogenetic analysis of selected full F-gene sequences was carried out using BEAST v 1.1.10.4 (Suchard et al. 2018) in combination with Beagle library (Ayres et al. 2012). We employed Uncorrelated relaxed clock and GMRF Bayesian SkyRide model (see Table S2), using General Time Reversal (GTR) substitution model was utilised, with separate partitions for codon positions 1 plus 2 versus position 3. Two independent MCMC chains with a length of 200,000,000 and sampling every 20,000 iterations were conducted, with the first 10% discarded as the burn-in. Convergence was assessed using Tracer v1.7.2 (Rambaut et al. 2018). The maximum clade credibility (MCC) tree was summarised using TreeAnnotator v1.10.4 (Rambaut et al. 2018) and visualized using R with the tidyverse (Wickham et al. 2019), treeio (L.-G. Wang et al. 2019) and phytools (Revell 2024) packages.

To determine regional spread, PPMV-1 samples were designated a region determined by the UN geoscheme (https://unstats.un.org/unsd/methodology/m49/), with the UK and Ireland designated independently as the British Isles. To explore the pattern of spatial diffusion among the geographic regions, discrete phylogeographic analyses using location as a trait were performed (Lemey et al. 2009). We assumed a symmetric non-reversible transition model and incorporated Bayesian stochastic search variable selection (Lemey et al. 2009). SpreaD3 (Bielejec et al. 2016) was used to measure rates of transmission using Bayes Factor (BF) and was used to determine likelihood of transmission between locations. The support for BF transmission was as described previously (Lee and Wagenmakers 2014). BF and representative transitions were visualised as described previously (Banyard et al. 2024). Markov jump counts were used to measure the number of viral movements along the branches of the phylogeny and estimated the Markov rewards to quantify the time the virus spent in each geographical region (Minin and Suchard 2008).

An alternative method to determine evolutionary history and geographical spread, analysis was determined using the mugration model in TreeTime with default settings (Sagulenko, Puller, and Neher 2018). Analysis was conducted on the M-L tree previously generated, with molecular clock estimation, again, utilising WHO regions as stated above.

## Results

### Phylogenetic relationship of the F-gene and complete genomes of PPMV-1 isolates from the British Isles

Following sequencing, the F-gene cleavage site was characterised as stated in Table 1. F-gene sequences previously designated as genotype VI, XX or XXI (PPMV-1-like) that were in the ND database were included in the analysis, along with isolates from the British Isles sequenced during this study. NCBI and BV-BRC databases were also analysed with sequences from Columbiformes included that were subsequently demonstrated to be PPMV-1-like. Two viruses isolated from poultry originating in the Middle East from 1968, Abu Ghraib 68 (AG68), Iraq, and Kuwait/256 (MK978147), were also included as it has been hypothesised that PPMV-1 originated in the Middle East during the 1960s, with the earliest detection of APMV-1 in pigeons being in Iraq in 1978 (Kaleta, Alexander, and Russell 1985); being previously described as lineage 4/Genotype VI-like. Two root samples were also included as suggested by Dimitrov *et al*., (2019). A Maximum-likelihood phylogeny of the British Isles PPMV-1 samples was produced and is shown in Figure **1**. All sequences analysed from Columbiformes detected in the British Isles were genotype VI or XXI, commonly associated with PPMV-1 (see Table 1). Analysis of the PPMV-1 F-gene sequences demonstrated that there have been three distinct dominant genotypes (i.e. more than a single genotype detection) in the British Isles since the first sequence from a virus in 1983.

**Figure 1:**
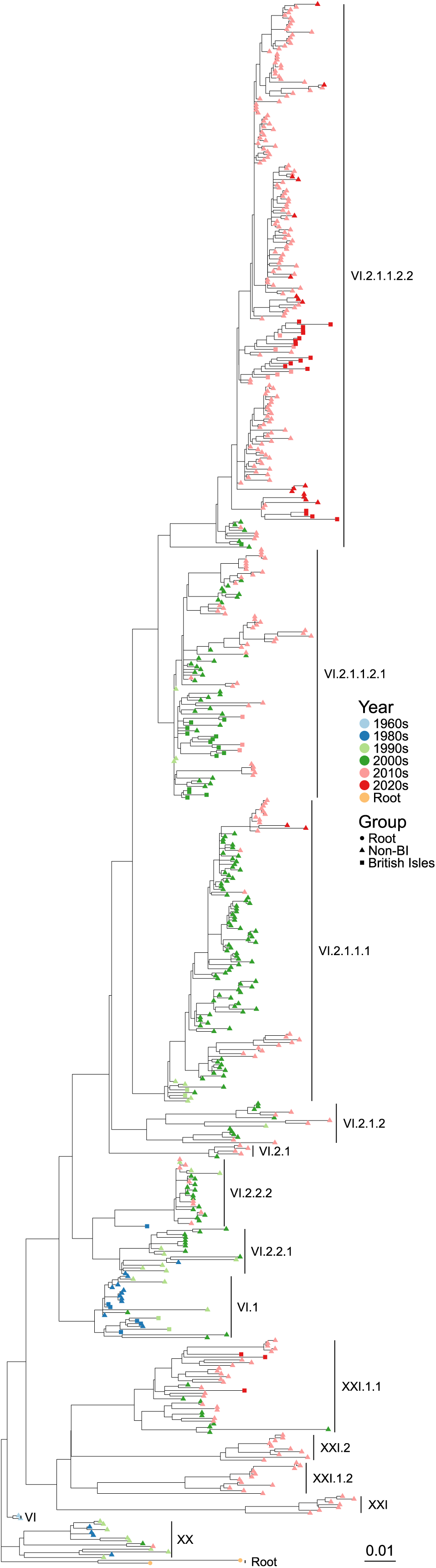
The British Isles have had multiple incursions of PPMV-1 from 1983 onwards. Maximum-likelihood tree of PPMV-1 F-gene sequences of APMV-1 isolates from Columbiformes or isolates previously shown to be PPMV-1-like. Genotypes are labelled as previously identified. Tips are coloured by decade of detection, and shape determines whether of British Isles origin or from elsewhere.

Viruses from 1983 - 1996 were of genotype VI.1 (previously called lineage 4b), commonly associated with viruses detected in Italy and the United States of America (USA), previously seen in the early 1980s, and appeared to be closely related to the first PPMV-1 detected in 1978 (Aldous et al. 2004). The 1989 virus (AV1528-89) was classified as a genotype VI.2.2.2, however, this was distinct from other VI.2.2.2 viruses analysed to-date, which were all previously detected in China. It suggests that the VI.2.2.2 viruses were transient throughout the British Isles and were not able to establish widespread infection in pigeons in the British Isles, or alternatively, the virus was not detected due to minimal surveillance activities. The 1997 virus (AV699-97) was a genotype VI.2.1.1.1 reported throughout Europe from 1996 (Ireland) through to 2000 (Italy) and was of the same genotype as a 1996 virus previously examined (GQ429292).

Between 2000 - 2011 viruses detected in samples from the British Isles were of the genotype VI.2.1.1.2.1, which was reported world-wide, with the first detection in Belgium in 1998 (accession numbers JX901109, JX901110 and JX901111). The first detection of genotype VI.1.2.2.1.1 in Columbiformes from the British Isles was in 2000 with the time to most recent common ancestor (TMRCA) estimated to be May 1996 (range October 1991 – November 1998 [TreeTime, August 1996; range November 1994 – July 1997]) (Figure 2, Box 1 and S1, Box 1). This isolate had no clear links to the initial viruses from Belgium but appeared to be more closely related to viruses from Italy, Luxembourg and Belgium detected from early 2000 onwards. Of the seventeen PPMV-1 VI.2.1.1.2.1 sequences generated here, a further three were phylogenetically linked to this initial incursion into the British Isles. Eight of the viruses were distinct from the initial incursion (Figure 2, box 2 and S1), with the TMRCA estimated to be June 1996 (range December 1992 – July 1999 [TreeTime, August 1996; range August 1995 – December 1997]), and this group included the last UK NDV isolate in Grey Partridges (*P. perdix*), which was hypothesised to be contracted from feral pigeons (Irvine et al. 2009). The remaining four isolates (from 2001, 2002 and 2004, Figure 2, Box 3 and S1, Box 3) were the most closely related to viruses detected in Belgium in 1998, with a TMRCA to those sequences of November 1997 (range January 1992 – November 1997 [TreeTime, September 1997; range March 1997 – January 1998]). The last genotype (VI.2.1.1.2.1) found in the UK and Ireland was in 2011 and was phylogenetically linked to the PPMV-1 isolates from the British Isles in 2000 (Figure 2, Box 1, Figure S1, Box1).

**Figure 2:**
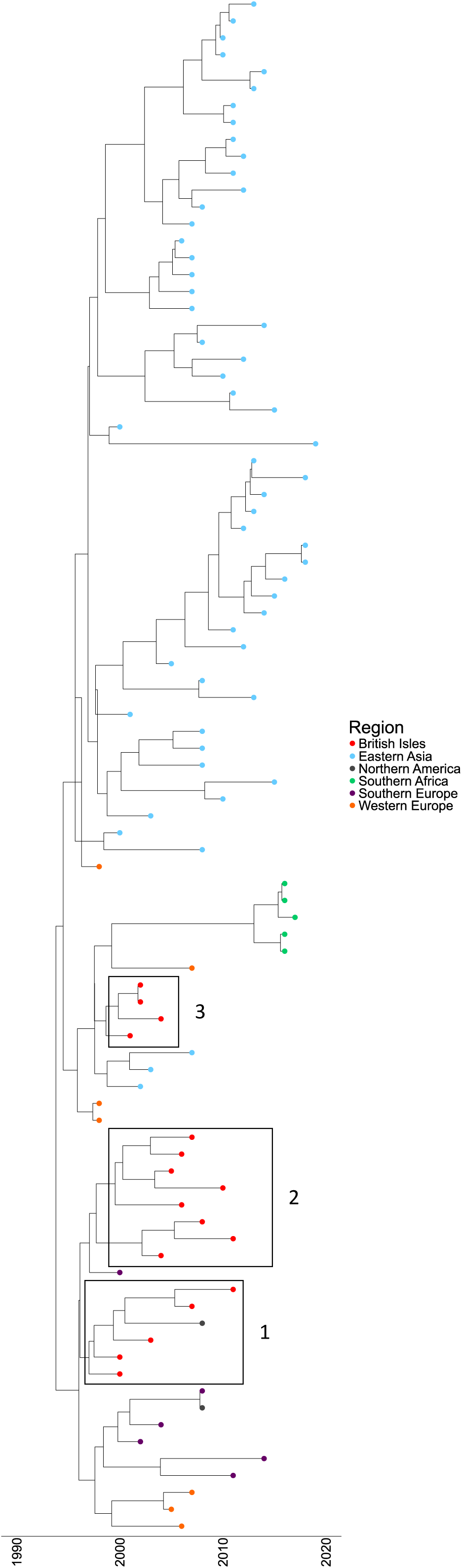
PPMV-1 genotype VI.2.1.1.2.1 incursion into the British Isles. Maximum clade credibility (MCC) phylogeny of PPMV-1 F-gene sequences identified as genotype VI.2.1.1.2.1. Phylogeny is scaled by year of collection. Tips are coloured by region of origin.

Between 2013 – 2023, 23 of the 26 characterised viruses were of the genotype VI.2.1.1.2.2, and all were closely related to viruses seen in Belgium in the early 2000s, and it appears that this is now the dominant, established PPMV-1 genotype within the British Isles. Within the three distinct genotype shifts, there were also novel genotypes seen present only in a single virus sequence analysed (Figure 1).

In 2021, the first identification of genotype XXI.1.1 PPMV-1 was observed within the British Isles. A second genotype XXI.1.1 was subsequently reported in the Channel Islands (Guernsey) in 2021, whilst a third was detected in 2022. This genotype was originally observed in Eastern Europe, but has since also been detected in the Ukraine, Egypt, Iran, and Pakistan amongst other countries (Sabra et al. 2017; Shabbir et al. 2024)

### Phylogenetic and geographic analysis of Genotype VI.2.1.1.2.2 PPMV-1 in the British Isles

Analysis of the PPMV-1 F-gene sequences demonstrated that the current genotype of PPMV-1 predominantly found in the British Isles was the VI.2.1.1.2.2 variant when analysed by Maximum-likelihood (M-L) phylogeny (Figure 1). We therefore focused on this genotype and determined the most recent common ancestors prior to arrival in the British Isles and determined the region from where initial incursions occurred. Locations of the twenty-three genotype VI.2.1.1.2.2 isolates identified were found throughout the British Isles are shown in Figure S2. The first identification of a genotype VI.2.1.1.2.2 worldwide was from a virus detected in Belgium in 2005, with the first detection in the British Isles occurring in November 2009 (Figure 1, 3A and S3). The TMRCA occurring with a Belgium virus detected from 2008 was predicted to have occurred in April 2005 (range November 2002 to April 2006) (Figure 3A) (TreeTime TMRCA estimate November 2005, range August 2004 – April 2007 Figure S3). Concurrently, the previously circulating VI.2.1.1.2.1 was also circulating worldwide, with the last British Isles detection in 2011. A virus from genotype VI.2.1.1.2.1 was also detected in Ireland in 2011, suggesting that this genotype was still the predominantly circulating PPMV-1 in Europe at this time.

**Figure 3:**
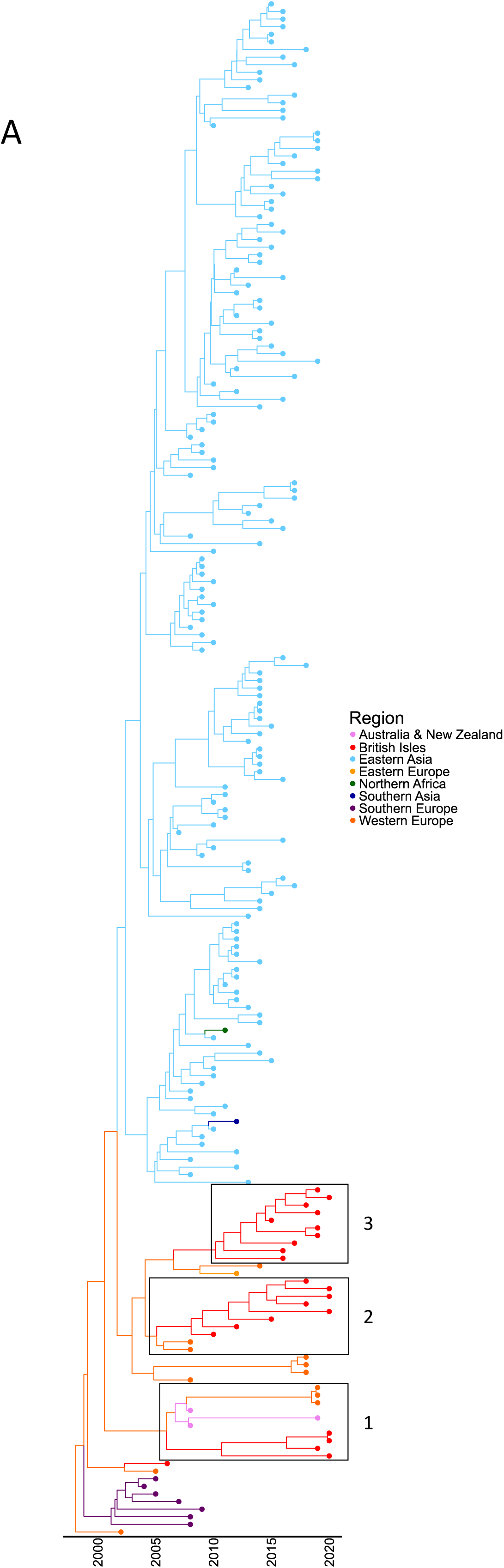
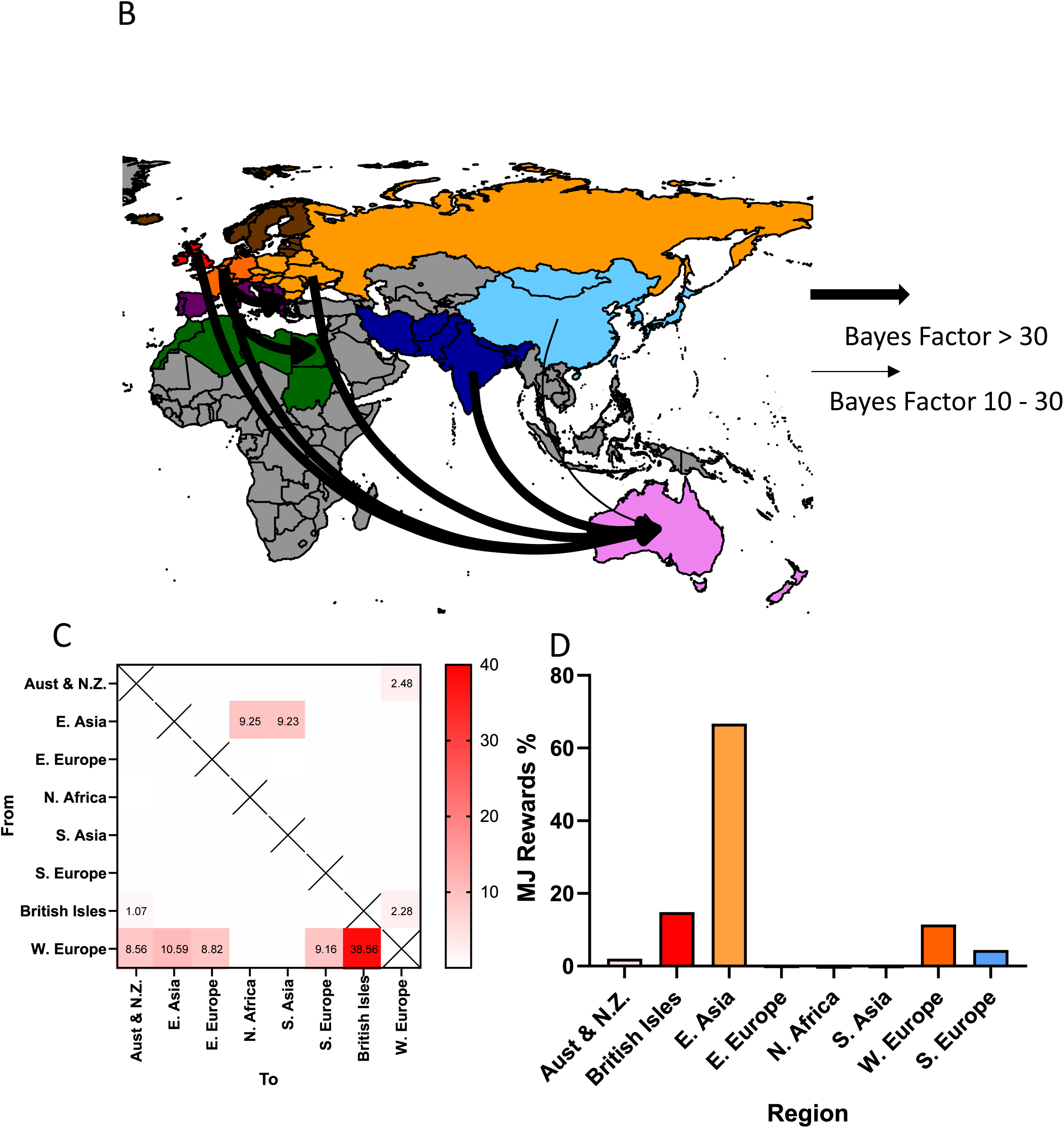
PPMV-1 genotype VI.2.1.1.2.2. incursion into the British Isles originated in Europe. **A)** MCC phylogeny of PPMV-1 F-gene sequences previously identified as genotype VI.2.1.1.2.2. Phylogeny is scaled to year of collection. Tips and branches are coloured by region of origin. **B)** Analysis of spread of PPMV-1 to different regions. Regions are coloured as per the UN geoscheme, with colours equivalent to regions identified in 3A. Thickness of arrow denotes strength of support. C) Frequency of transitions determined by Markov jump counts genotype VI.2.1.1.2.2 isolates between different regions, D) Proportion of time the virus spent in each region.

Genetic analysis suggests that there have been a further three independent introductions of genotype VI.2.1.1.2.2 isolates into the British Isles after the initial incursion in 2009, with all three phylogenetically distinct from the initial incursion (Figure 3A and S3). Descendants of three incursions are still circulating within the British Isles, sharing common ancestors from mainland Europe. The initial viruses were detected in June 2013 (Figure 3A, Box 2 and S3, Box 2) and were most closely related to PPMV-1 viruses detected in Belgium in 2011 (TMRCA February 2008, range November 2005 – November 2009, TreeTime TMRCA estimate October 2008 range May 2008 – September 2009). The second incursion was detected in a pigeon from October 2018 (Figure 3A, Box 3, S3 Box 3) and was most closely related to viruses from Ukraine from 2015 (TMRCA July 2009, range August 2006 – April 2012, TreeTime TMRCA May 2009, range July 2008 – October 2010). The third incursion was first detected in the British Isles in September 2022 (Figure 3A Box 1, S3 Box 1), with the most closely virus being from Australia in 2011 (TMRCA December 2008, range August 2006 – August 2010, TreeTime TMRCA estimate June 2010, range December 2009 to November 2011). The derivatives of these three independent genotype VI.2.1.1.2.2 incursions are still circulating concurrently within the British Isles in 2023 (when the period for this analysis was concluded).

Genotype VI.2.1.1.2.2 was initially detected in Europe in 2006, with the first non-European VI.2.1.1.2.2 detected in China in 2010. The estimated time of the last common European ancestor prior to incursion into China by MCC analysis was August 2004 (range October 2002 to May 2006) (TreeTime TMRCA estimate February 2007, range August 2006 – April 2008 (Figure S3)), so there appears to be a significant gap between initial incursion and the primary detection of PPMV-1 genotype VI.2.1.1.2.2 in non-European countries. Genotype VI.2.1.1.2.2 now appears to be the predominant PPMV-1 genotype in China, with the last detection of VI.2.1.1.2.1 occurring in December 2018 (MZ306219) and VI.2.2.2 in 2013 (KT381596). The Markov jump analysis shows a strong correlation with movement from Europe to Asia. Movement of PPMV-1 genotype VI.2.1.1.2.2 from Europe to Asia could have occurred through direct interaction of Columbiformes, with onward transmission to Asia via several unsampled intermediates, or through anthropogenic movement. Sporadic detections in other continents are seen, however it does appear, with the data available, that genotype VI.2.1.1.2.2 is the common PPMV-1 detected throughout Europe and China. However, there are distinct European and Asian lineages of genotype VI.2.1.1.2.2, with clear monophyly for each region.

PPMV-1 genotype VI.2.1.1.2.2 was also detected in Australia in 2011 with this isolate being of European origin with a TMRCA estimate of November 2009 (range December 2007 – October 2010, TreeTime TMRCA estimate April 2010 range September 2009 – October 2010) (Figures 3A and S3)). Viruses phylogenetically linked to the Australian PPMV-1s were subsequently observed back in Europe, with these detections occurring in Switzerland in 2022, and appearing in the British Isles in 2023, but this does appear to be infrequent.

To determine potential routes of transmission, Bayesian stochastic search variable selections (BSSVS) were conducted, using the regions defined by the UN geoscheme from which the sequences originated as a discrete trait. Support for transmission was quantified by Bayes Factor (BF). This analysis demonstrated very strong BF (> 30) for transmission to Australia and New Zealand (NZ) from the UK and Ireland, Western Europe and Southern Asia, and from Western Europe to both Southern Europe and Northern Africa (See Figure 3B and table S3). Very strong BF (30 – 100) was observed for transmission from Eastern Europe to Australia and NZ, whilst strong BF (10 – 30) was observed from Eastern Asia to Australia and NZ. Analysis using BF does not determine the routes of transmission into the British Isles for genotype VI.2.1.1.2.2 viruses, or the direct movement of PPMV-1 from Australia and NZ back into Europe.

Markov jump analysis suggests that the genotype VI.2.1.1.2.2 isolates have a high percentage of transition from Western Europe into other regions examined, with the highest number of transitions observed from Western Europe to the British Isles, with fewer predicted transitions in the opposite direction (Figure 3C). Transitions from Western Europe to Eastern and Southern Europe were also predicted to have occurred, whilst transition from Western Europe to Eastern Asia were also predicted, with minimal evidence of movement back to Europe (Figure 3C), in agreement with previous analysis (Xie et al. 2020). There were, however, predicted transitions from Western Europe to Australia/NZ, alongside transitions occurring from British Isles to Australia/NZ. However, there were predicted transitions from Australia/NZ back to both Western Europe. Isolates observed in North Africa and Southern Asia were likely to have transitioned from Eastern Asia. From this analysis and the current data available, genotype VI.2.1.1.2.2 originated in Western Europe, before spreading through Europe, Asia and Australia. Analysis of Markov rewards with the available data suggests that currently, the genotype VI.2.1.1.2.2 isolates have spent the longest time in Eastern Asia, followed closely by British Isles and Western Europe (Figure 3D). The spread and Markov jump analysis are dependent on sequences currently available in the database. Due to the paucity of PPMV-1 sequences, it is difficult to determine the accuracy of these predictions, with some regions under- or over-sampled, which may bias analysis.

## Discussion

The emergence, maintenance and detection of avian paramyxoviruses in pigeons and doves is poorly defined, primarily due to the varied, dynamic and feral distribution of these species globally. Many pigeon species are non-migratory and so opportunity for global spread is limited to displacement by anthropogenic factors or ecological dispersal mechanisms. Therefore, the opportunity for genotype distribution between populations must be considered to be low and viral diversity should be limited. However, PPMV-1 is endemic in Columbiformes globally and has been detected in pigeons in the Middle East since the 1960s. Following the initial identification in pigeons in 1978 (Kaleta, Alexander, and Russell 1985), it was subsequently identified throughout Europe in the 1980s, with the first detection in the British Isles in 1983 (Lister, Alexander, and Hogg 1986).

Previous studies have examined the genetic diversity of PPMV-1 in the UK including the assessment of 94 PPMV-1 sequences between 1978 - 2002 (Aldous et al. 2004) and subsequently analysis involving a further 43 PPMV-1 sequences from 2003 – 2010. The current study has enhanced this data set through the generation of sequence data for a further 51 PPMV-1 detections from across the British Isles including 39 sequences generated using the Illumina platform, eight via Sanger sequencing, and four sequences generated F-gene nanopore sequencing, with isolates ranging from 1983 - 2023. Analysis of samples ranging from 1983 – 2010 demonstrated that the genotypes detected matched with the previously defined lineages/genotypes circulating in the British Isles described by previous studies (Aldous et al. 2010; 2004). From these analyses, pigeon or dove species appear to have maintained three distinct PPMV-1 genotypes since the initial detection in 1983. Those initial detections were genotype VI.1 (lineage 4b, 4bi, 4c, genotype VIb) and included viruses from 1983 – 1996 (Aldous et al. 2004). Genotype VI.2.1.1.2.1 (lineage 4b*ii* 4f, genotype Vij), was then detected in 1999 in the UK, a year prior to the first sequence analysed in this study (Aldous et al. 2004). This, however, does not predate the initial detection of VI.2.1.1.2.1 viruses from Belgium and Portugal in 1998 (Aldous et al. 2010), which again suggests that incursion into the British Isles was likely from mainland Europe. The third genotype incursion into the British Isles was VI.2.1.1.2.2, again with the earliest detection observed in Belgium in 2005 (Van Borm et al. 2013) with the first British Isles detection in 2009, and this genotype appearing to become the dominant PPMV-1 genotype across Europe and China from this point onwards (Zhan et al. 2022; Tian et al. 2020; Van Borm et al. 2013; Annaheim et al. 2022). Sporadic genotypes were also observed in the British Isles throughout this period of study, specifically a VI.2.2.2 in 1989 and VI.2.1.1 in1997. Interestingly, three genotype XXI.1.1 (previously lineage 4a/d, genotype VIg) viruses were detected from 2021 onwards. This genotype was first observed in Russia in 2005 (Pchelkina et al. 2013) and now appears to be increasing in prevalence across Southern Asia, Africa, and Eastern Europe (Sabra et al. 2017).

The movement of PPMV-1 from mainland Europe to the BIs is not an unexpected occurrence due to closeness of land masses which may allow for feral Columbiformes to interact. Coupled with that, and the frequence interactions of both racing pigeons and show birds throughout the BIs and Europe, has the potential for onward infection and spread from one region to another. The subsequent spread of genotype VI.2.1.1.2.2 from the initial detection in Europe through to Asia and into Australia occurred at separate times and through different common ancestors. The mechanism of spread through these regions is difficult to determine, especially from Europe to Australia and back. Pigeons and doves are considered not to migrate, with birds staying close to their birth site. Rock Doves and feral pigeons, common in many cities worldwide, are thought to travel as little as 0.5 km per day (Rose, Nagel, and Haag-Wackernagel 2006). Long distance migration of Columbiformes is thought to be uncommon, although the Common Wood Pigeon migrate from Scandinavia to Southern Spain (Schumm et al. 2022). The limited sequencing information makes it difficult to precisely determine the mechanism of incursion into Australia, although, with no clear linkage to PPMV-1 samples from South-East Asia, and no obvious migration route by Columbiformes, the most parsimonious hypothesis is due to the human import of infected pigeons or materials into this region. How this virus returned is also unclear, although again, the role of humans in this process cannot be discounted. However, determining the time of introduction initially into Australia, and then the re-introduction into Europe is difficult due to substantial unsampled ancestry and the lack of intermediates PPMV-1 sequences, a common issue when analysing transmission of PPMV-1 from region to region. The increased trade in racing and show pigeons between different regions may also increase transmission of novel PPMV-1 isolates, along with other pigeon pathogens, and the potential import of novel pigeon pathogens to different continents need to be considered when trade occurs.

Analysis of TMRCA using two distinct methods, BEAST and Treetime, revealed differences in proposed TMRCA values and the associated confidence ranges. This is to be expected due to the different methods used to determine the TMRCA, however, the large branch-lengths (possibly due to unsampled ancestry) observed for some of this analysis will make TMRCA analysis difficult and reduce the accuracy of an estimation. Analysis to further understand spread of genotype VI.2.1.1.2.2 was undertaken, with limited identification of spread throughout large regions of the world, with only introductions into Australia, North Africa and Eastern Europe identified. Indeed, there is a distinct issue with the analysis of PPMV-1, with limited sequencing and general paucity of surveillance initiatives which has inhibited the understanding of the spread of PPMV-1 globally. The lack of sampling and virus definition may impact upon undefined increases in virulence determinants for other species including poultry as well as the risk of re-emergence of novel PPMV-1 isolates capable of infecting poultry. It is therefore of paramount importance to increase the sampling of Columbiformes, both captive and feral to ascertain the potential risk more precisely to poultry production. Indeed, an increase in detection of ND in European countries linked to contemporary PPMV-1 isolates has occurred over since 2022 (Annaheim et al. 2022; AI-ND EURL 2024; 2023), demonstrating the importance of continual surveillance for PPMV-1.

PPMV-1 is endemic in Columbiformes world-wide, and thus, the evolutionary pressures involved in adaptation to this family of birds has resulted in genetic changes which may confer continued transmission as opposed to other avian species. Due to the presence of a virulent F-gene cleavage site, the detection of PPMV-1 in poultry would result in the declaration of an ND outbreak as per the WOAH definition (World Organisation for Animal Health (WOAH) 2021). However, recent analysis of the genotype VI.2.1.1.2.2 isolates, which are the most contemporary identified within the UK, and across China and Europe, suggests that infection of poultry with these viruses does not result in definitive clinical signs and reduced mortality compared to PPMV-1 from the 1980s (Zhan et al. 2022), whilst infection of pigeons does appear to show low-level morbidity and mortality in comparison (Zhan et al. 2022; Yu et al. 2022; Tian et al. 2020). However, with the continued presence of PPMV-1 in a substantial population of wild birds, the need for increased surveillance to aid risk definition and compliance with good biosecurity for poultry, with distinct separation of poultry from wild birds and Columbiformes essential to ensure that outbreaks of ND are kept to a minimum.

## Acknowledgements

We wish to thank Sahar Mahmood and Paul Skinner for virus propagation and sequence analysis along with the members of the WOAH international reference laboratory for providing isolates. We also wish to thank the Central Unit Sequencing and PCR (CUSP) at APHA Weybridge for generating whole genome sequences.

---

**Figure S1 PPMV-1 genotype VI.2.1.1.2.1 incursion into the British Isles.** Treetime analysis of M-L phylogeny of PPMV-1 F-gene sequences previously identified as genotype VI.2.1.1.2.1. Phylogeny is scaled by year of collection. Tips are coloured by region of origin.

**Figure S2 Locations of PPMV-1 genotype VI.2.1.1.2.2 in the United Kingdom.** The locations of the twenty-three genotype VI.2.1.1.2.2 isolates identified for this study. Sites are coloured according to the decade that they were found.

**Figure S3 PPMV-1 genotype VI.2.1.1.2.2 incursion into the British Isles**. Treetime and mugration analysis of M-L phylogeny of isolates identified as genotype VI.2.1.1.2.2. Phylogeny is determined by year of collection. Tips and branches are coloured by region of origin.

## References

AI-ND EURL. 2019. “MEMBER STATE (& THIRD COUNTRIES) REPORTS FOR AVIAN AVULAVIRUSES 2019.” 2019. https://www.izsvenezie.com/documents/reference-laboratories/avian-influenza/workshops/2020/23-member-state-thid-countries-reports-for-avian-orthoavulaviruses-2019.pdf.

AI-ND EURL. 2023. “29th Annual Meeting of the National Reference Laboratories for Avian Influenza and Newcastle Disease of European Union Member States (2023 – Parma, Italy).” 2023. https://www.izsvenezie.com/reference-laboratories/avian-influenza-newcastle-disease/workshops/.

AI-ND EURL. 2024. “30th Annual Meeting of the National Reference Laboratories for Avian Influenza and Newcastle Disease of European Union Member States (2024 – Venice Mestre, Italy).” 2024. https://www.izsvenezie.com/reference-laboratories/avian-influenza-newcastle-disease/workshops/.

Aldous, E. W., C. M. Fuller, J. H. Ridgeon, R. M. Irvine, D. J. Alexander, and I. H. Brown. 2014. “The Evolution of Pigeon Paramyxovirus Type 1 (PPMV-1) in Great Britain: A Molecular Epidemiological Study.” Transboundary and Emerging Diseases 61 (2): 134–39. 10.1111/tbed.12006.

Aldous, E. W., J. K. Mynn, J. Banks, and D. J. Alexander. 2003. “A Molecular Epidemiological Study of Avian Paramyxovirus Type 1 (Newcastle Disease Virus) Isolates by Phylogenetic Analysis of a Partial Nucleotide Sequence of the Fusion Protein Gene.” Avian Pathol 32 (3): 239–56. 10.1080/030794503100009783.

Aldous, E. W., J. K. Mynn, R. M. Irvine, D. J. Alexander, and I. H. Brown. 2010. “A Molecular Epidemiological Investigation of Avian Paramyxovirus Type 1 Viruses Isolated from Game Birds of the Order Galliformes.” Avian Pathology 39 (6): 519–24. 10.1080/03079457.2010.530938.

Aldous, E W, C M Fuller, J K Mynn, and D J Alexander. 2004. “A Molecular Epidemiological Investigation of Isolates of the Variant Avian Paramyxovirus Type 1 Virus (PPMV-1) Responsible for the 1978 to Present Panzootic in Pigeons.” Avian Pathol 33 (2): 258–69. 10.1080/0307945042000195768.

Alexander, D. J., R. J. Manvell, K. M. Frost, W. J. Pollitt, D. Welchman, and K. Perry. 1997. “Newcastle Disease Outbreak in Pheasants in Great Britain in May 1996.” Veterinary Record 140 (1): 20–22. 10.1136/vr.140.1.20.

Alexander, D. J., G. Parsons, and R. Marshall. 1984. “Infection of Fowls with Newcastle Disease Virus by Food Contaminated with Pigeon Faeces.” The Veterinary Record 115 (23): 601–2. 10.1136/vr.115.23.601.

Alexander, D J, P H Russell, and M S Collins. 1984. “Paramyxovirus Type 1 Infections of Racing Pigeons: 1 Characterisation of Isolated Viruses.” The Veterinary Record 114 (18): 444–46. 10.1136/vr.114.18.444.

Alexander, Dennis J. 2011. “Newcastle Disease in the European Union 2000 to 2009.” Avian Pathology 40 (6): 547–58. 10.1080/03079457.2011.618823.

Annaheim, Désirée, Barbara Renate Vogler, Brigitte Sigrist, Andrea Vögtlin, Daniela Hüssy, Christian Breitler, Sonja Hartnack, et al. 2022. “Screening of Healthy Feral Pigeons (Columba Livia Domestica) in the City of Zurich Reveals Continuous Circulation of Pigeon Paramyxovirus-1 and a Serious Threat of Transmission to Domestic Poultry.” Microorganisms 10 (8). 10.3390/microorganisms10081656.

Ayres, Daniel L., Aaron Darling, Derrick J. Zwickl, Peter Beerli, Mark T. Holder, Paul O. Lewis, John P. Huelsenbeck, et al. 2012. “BEAGLE: An Application Programming Interface and High-Performance Computing Library for Statistical Phylogenetics.” Systematic Biology 61 (1): 170–73. 10.1093/sysbio/syr100.

Banyard, Ashley C, Ashley Bennison, Alexander M P Byrne, Scott M Reid, Joshua G Lynton-Jenkins, Benjamin Mollett, Dilhani De Silva, et al. 2024. “Detection and Spread of High Pathogenicity Avian Influenza Virus H5N1 in the Antarctic Region.” Nature Communications 15 (1): 7433. 10.1038/s41467-024-51490-8.

Biancifiori, F, and A Fioroni. 1983. “An Occurrence of Newcastle Disease in Pigeons: Virological and Serological Studies on the Isolates.” Comp Immunol Microbiol Infect Dis 6 (3): 247–52. 10.1016/0147-9571(83)90017-6.

Bielejec, Filip, Guy Baele, Bram Vrancken, Marc A. Suchard, Andrew Rambaut, and Philippe Lemey. 2016. “SpreaD3: Interactive Visualization of Spatiotemporal History and Trait Evolutionary Processes.” Molecular Biology and Evolution 33 (8): 2167–69. 10.1093/molbev/msw082.

Borm, Steven Van, Toon Rosseel, Mieke Steensels, Thierry Van den Berg, and Bénédicte Lambrecht. 2013. “What’s in a Strain? Viral Metagenomics Identifies Genetic Variation and Contaminating Circoviruses in Laboratory Isolates of Pigeon Paramyxovirus Type 1.” Virus Research 171 (1): 186–93. 10.1016/j.virusres.2012.11.017.

Byrne, Alexander M P, Joe James, Benjamin C Mollett, Stephanie M Meyer, Thomas Lewis, Magdalena Czepiel, Amanda H Seekings, et al. 2023. “Investigating the Genetic Diversity of H5 Avian Influenza Viruses in the United Kingdom from 2020-2022.” Microbiology Spectrum 11 (4): e0477622. 10.1128/spectrum.04776-22.

Collins, M. S., J. B. Bashiruddin, and D. J. Alexander. 1993. “Deduced Amino Acid Sequences at the Fusion Protein Cleavage Site of Newcastle Disease Viruses Showing Variation in Antigenicity and Pathogenicity.” Archives of Virology 128 (3–4): 363–70. 10.1007/BF01309446.

Department for Environment Food and Rural Affairs. 2019. Notifiable Avian Disease Control Strategy for Great Britain. https://assets.publishing.service.gov.uk/media/5d8cc8ceed915d556efe16cf/avian-disease-control-strategy1.pdf.

Diel, Diego G., Luciana H.A. da Silva, Hualei Liu, Zhiliang Wang, Patti J. Miller, and Claudio L. Afonso. 2012. “Genetic Diversity of Avian Paramyxovirus Type 1: Proposal for a Unified Nomenclature and Classification System of Newcastle Disease Virus Genotypes.” Infection, Genetics and Evolution 12 (8): 1770–79. 10.1016/j.meegid.2012.07.012.

Dimitrov, K M, C Abolnik, C L Afonso, E Albina, J Bahl, M Berg, F X Briand, et al. 2019. “Updated Unified Phylogenetic Classification System and Revised Nomenclature for Newcastle Disease Virus.” Infection, Genetics and Evolution 74:103917. 10.1016/j.meegid.2019.103917.

Dimitrov, K M, A M Ramey, X Qiu, J Bahl, and C L Afonso. 2016. “Temporal, Geographic, and Host Distribution of Avian Paramyxovirus 1 (Newcastle Disease Virus).” Infect Genet Evol 39:22–34. 10.1016/j.meegid.2016.01.008.

Gibbs, David. 2010. Pigeons and Doves: A Guide to the Pigeons and Doves of the World. London: Bloomsbury Publishing.

Graziosi, Giulia, Caterina Lupini, Francesco Dalla Favera, Gabriella Martini, Geremia Dosa, Giacomo Trevisani, Gloria Garavini, Alessandro Mannelli, and Elena Catelli. 2024. “Characterizing the Domestic-Wild Bird Interface through Camera Traps in an Area at Risk for Avian Influenza Introduction in Northern Italy.” Poultry Science 103 (8): 103892. 10.1016/j.psj.2024.103892.

Irvine, R. M., E. W. Aldous, R. J. Manvell, W. J. Cox, V. Ceeraz, C. M. Fuller, A. M. Wood, et al. 2009. “Outbreak of Newcastle Disease Due to Pigeon Paramyxovirus Type 1 in Grey Partridges (Perdix Perdix) in Scotland in October 2006.” The Veterinary Record 165 (18): 531–35. 10.1136/vr.165.18.531.

Kaleta, E F, D J Alexander, and P H Russell. 1985. “The First Isolation of the Avian PMV-1 Virus Responsible for the Current Panzootic in Pigeons ?” Avian Pathol 14 (4): 553–57. 10.1080/03079458508436258.

Lee, Michael D, and Eric-Jan Wagenmakers. 2014. “Bayesian Model Comparison.” In Bayesian Cognitive Modeling: A Practical Course, 101–117. Cambridge University Press.

Lemey, Philippe, Andrew Rambaut, Alexei J. Drummond, and Marc A. Suchard. 2009. “Bayesian Phylogeography Finds Its Roots.” PLoS Computational Biology 5 (9): 1000520. 10.1371/journal.pcbi.1000520.

Lister, S. A., D. J. Alexander, and R. A. Hogg. 1986. “Evidence for the Presence of Avian Paramyxovirus Type 1 in Feral Pigeons in England and Wales.” The Veterinary Record 118 (17): 476–79. 10.1136/vr.118.17.476.

Meulemans, G, T P van den Berg, M Decaesstecker, and M Boschmans. 2002. “Evolution of Pigeon Newcastle Disease Virus Strains.” Avian Pathol 31 (5): 515–19. 10.1080/0307945021000005897.

Minin, Vladimir N., and Marc A. Suchard. 2008. “Fast, Accurate and Simulation-Free Stochastic Mapping.” Philosophical Transactions of the Royal Society B: Biological Sciences 363 (1512): 3985–95. 10.1098/rstb.2008.0176.

Pchelkina, I. P., T. B. Manin, S. N. Kolosov, S. K. Starov, A. V. Andriyasov, I. A. Chvala, V. V. Drygin, Q. Yu, P. J. Miller, and D. L. Suarez. 2013. “Characteristics of Pigeon Paramyxovirus Serotype-1 Isolates (PPMV-1) from the Russian Federation from 2001 to 2009.” Avian Diseases 57 (1): 2–7. 10.1637/10246-051112-Reg.1.

Rambaut, Andrew, Alexei J. Drummond, Dong Xie, Guy Baele, and Marc A. Suchard. 2018. “Posterior Summarization in Bayesian Phylogenetics Using Tracer 1.7.” Systematic Biology 67 (5): 901–4. 10.1093/sysbio/syy032.

Reid, Scott M., Paul Skinner, David Sutton, Craig S. Ross, Karolina Drewek, Natalia Weremczuk, Ashley C. Banyard, et al. 2023. “Understanding the Disease and Economic Impact of Avirulent Avian Paramyxovirus Type 1 (APMV-1) Infection in Great Britain.” Epidemiology and Infection 151:e163. 10.1017/S0950268823001255.

Revell, Liam J. 2024. “Phytools 2.0: An Updated R Ecosystem for Phylogenetic Comparative Methods (and Other Things).” PeerJ 12:e16505. 10.7717/peerj.16505.

Rose, Eva, Peter Nagel, and Daniel Haag-Wackernagel. 2006. “Spatio-Temporal Use of the Urban Habitat by Feral Pigeons (Columba Livia).” Behavioral Ecology and Sociobiology 60:242–254. 10.1007/s00265-006-0162-8.

Sabra, Mahmoud, Kiril M. Dimitrov, Iryna V. Goraichuk, Abdul Wajid, Poonam Sharma, Dawn Williams-Coplin, Asma Basharat, et al. 2017. “Phylogenetic Assessment Reveals Continuous Evolution and Circulation of Pigeon-Derived Virulent Avian Avulaviruses 1 in Eastern Europe, Asia, and Africa.” BMC Veterinary Research 13 (1): 291. 10.1186/s12917-017-1211-4.

Sagulenko, Pavel, Vadim Puller, and Richard A. Neher. 2018. “TreeTime: Maximum-Likelihood Phylodynamic Analysis.” Virus Evolution 4 (1): vex042. 10.1093/ve/vex042.

Schumm, Yvonne R., Juan F. Masello, Valerie Cohou, Philippe Mourguiart, Benjamin Metzger, Sascha Rösner, and Petra Quillfeldt. 2022. “Should I Stay or Should I Fly? Migration Phenology, Individual-Based Migration Decision and Seasonal Changes in Foraging Behaviour of Common Woodpigeons.” Science of Nature 109 (5): 44. 10.1007/s00114-022-01812-x.

Shabbir, Muhammad Zubair, Sahar Mahmood, Aziz Ul-Rahman, Ashley C Banyard, and Craig S Ross. 2024. “Genomic Diversity and Evolutionary Insights of Avian Paramyxovirus-1 in Avian Populations in Pakistan.” Viruses 16 (9): 1414. 10.3390/v16091414.

Shan, Songhua, Kerri Bruce, Vittoria Stevens, Frank Y K Wong, Jianning Wang, Dayna Johnson, Deborah Middleton, et al. 2021. “In Vitro and In Vivo Characterization of a Pigeon Paramyxovirus Type 1 Isolated from Domestic Pigeons in Victoria, Australia 2011.” Viruses 13 (3): 429. 10.3390/v13030429.

Suchard, Marc A, Philippe Lemey, Guy Baele, Daniel L Ayres, Alexei J Drummond, and Andrew Rambaut. 2018. “Bayesian Phylogenetic and Phylodynamic Data Integration Using BEAST 1.10.” Virus Evolution 4 (1): vey016. 10.1093/ve/vey016.

Sutton, David A., David P. Allen, Chad M. Fuller, Jo Mayers, Benjamin C. Mollett, Brandon Z. Londt, Scott M. Reid, Karen L. Mansfield, and Ian H. Brown. 2019. “Development of an Avian Avulavirus 1 (AAvV-1) L-Gene Real-Time RT-PCR Assay Using Minor Groove Binding Probes for Application as a Routine Diagnostic Tool.” J Virol Methods 265:9–14. 10.1016/j.jviromet.2018.12.001.

Tian, Ye, Ruixue Xue, Wanting Yang, Yujie Li, Jia Xue, and Guozhong Zhang. 2020. “Characterization of Ten Paramyxovirus Type 1 Viruses Isolated from Pigeons in China during 1996–2019.” Veterinary Microbiology 244:108661. 10.1016/j.vetmic.2020.108661.

Wang, Li-Gen, Tommy Tsan-Yuk Lam, Shuangbin Xu, Zehan Dai, Lang Zhou, Tingze Feng, Pingfan Guo, et al. 2019. “Treeio: An R Package for Phylogenetic Tree Input and Output with Richly Annotated and Associated Data.” Molecular Biology and Evolution 37 (2): 599–603. 10.1093/molbev/msz240.

Wang, Zeren, Zhengyang Geng, Hongbo Zhou, Pengju Chen, Jing Qian, and Aizhen Guo. 2024. “Genetic Characterization, Pathogenicity, and Epidemiology Analysis of Three Sub-Genotype Pigeon Newcastle Disease Virus Strains in China.” Microorganisms 12 (4): 738. 10.3390/microorganisms12040738.

Wickham, Hadley, Mara Averick, Jennifer Bryan, Winston Chang, Lucy McGowan, Romain François, Garrett Grolemund, et al. 2019. “Welcome to the Tidyverse.” Journal of Open Source Software 4 (43). 10.21105/joss.01686.

World Organisation for Animal Health (WOAH). 2021. “Terrestrial Manual: Newcastle Disease (Infection with Newcastle Disease Virus).” 2021. https://www.woah.org/fileadmin/Home/eng/Health_standards/tahm/3.03.14_NEWCASTLE_DIS.pdf.

Xie, Peng, Libin Chen, Yifan Zhang, Qiuyan Lin, Chan Ding, Ming Liao, Chenggang Xu, Bin Xiang, and Tao Ren. 2020. “Evolutionary Dynamics and Age-Dependent Pathogenesis of Sub-Genotype VI.2.1.1.2.2 PPMV-1 in Pigeons.” Viruses 12 (4): 433. 10.3390/v12040433.

Yu, Xiaohui, Yaoyao Luo, Jingjing Wang, Bo Shu, Wenming Jiang, Shuo Liu, Yang Li, et al. 2022. “A Molecular, Epidemiological and Pathogenicity Analysis of Pigeon Paramyxovirus Type 1 Viruses Isolated from Live Bird Markets in China in 2014–2021.” Virus Research 318 (September):198846. 10.1016/j.virusres.2022.198846.

Zhan, Tiansong, Dongchang He, Xiaolong Lu, Tianxing Liao, Wenli Wang, Qing Chen, Xiaowen Liu, et al. 2021. “Biological Characterization and Evolutionary Dynamics of Pigeon Paramyxovirus Type 1 in China.” Frontiers in Veterinary Science 8:721102. 10.3389/fvets.2021.721102.

Zhan, Tiansong, Xiaolong Lu, Dongchang He, Xiaomin Gao, Yu Chen, Zenglei Hu, Xiaoquan Wang, Shunlin Hu, and Xiufan Liu. 2022. “Phylogenetic Analysis and Pathogenicity Assessment of Pigeon Paramyxovirus Type 1 Circulating in China during 2007–2019.” Transboundary and Emerging Diseases 69 (4): 2076–88. 10.1111/tbed.14215.

