## Supplementary Figures for "Phylogenetic analysis of pigeon paramyxovirus type 1 (PPMV-1) detected in the British Isles between 1983 – 2023": Supplementary Figures - Byrne et al.pdf

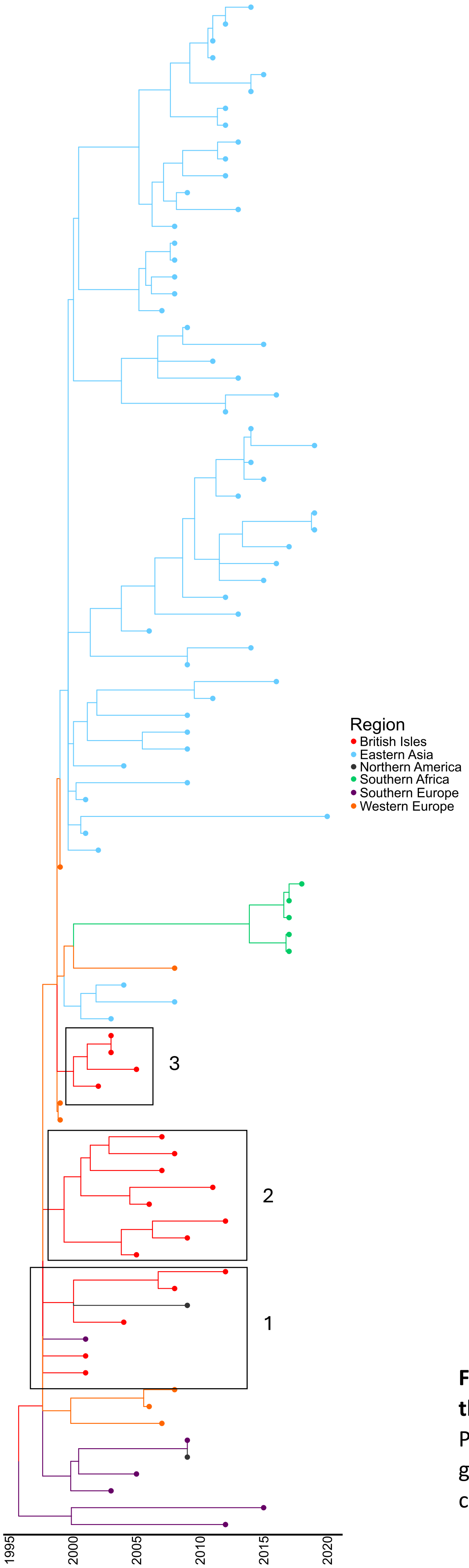


**Figure S1 PPMV-1 genotype VI.2.1.1.2.1 incursion into the British Isles.** Treetime analysis of M-L phylogeny of PPMV-1 F-gene sequences previously identified as genotype VI.2.1.1.2.1. Phylogeny is scaled by year of collection. Tips are coloured by region of origin.

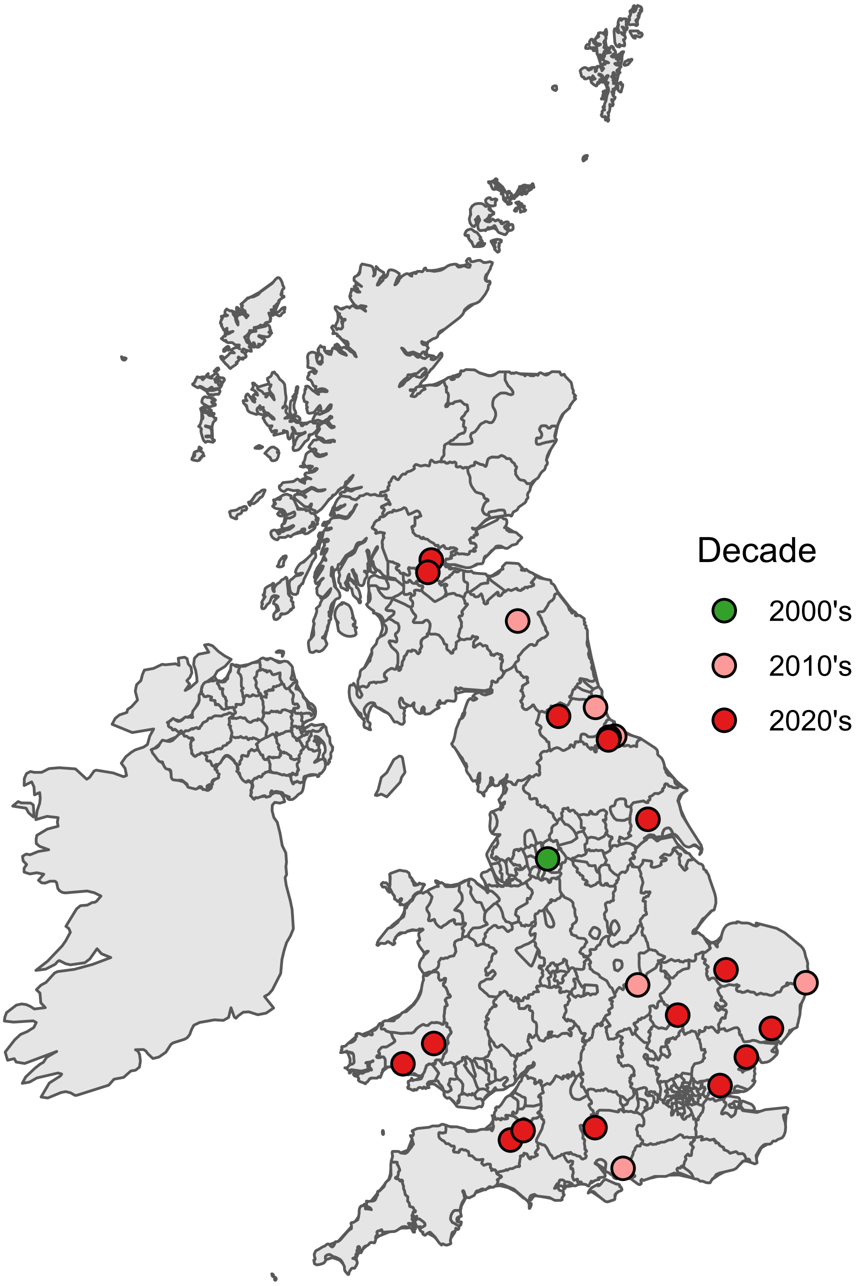


**Figure S2 Locations of PPMV-1 genotype VI.2.1.1.2.2 in the United Kingdom.** The locations of the twenty-three genotype VI.2.1.1.2.2 isolates identified for this study. Sites are coloured according to the decade that they were found.

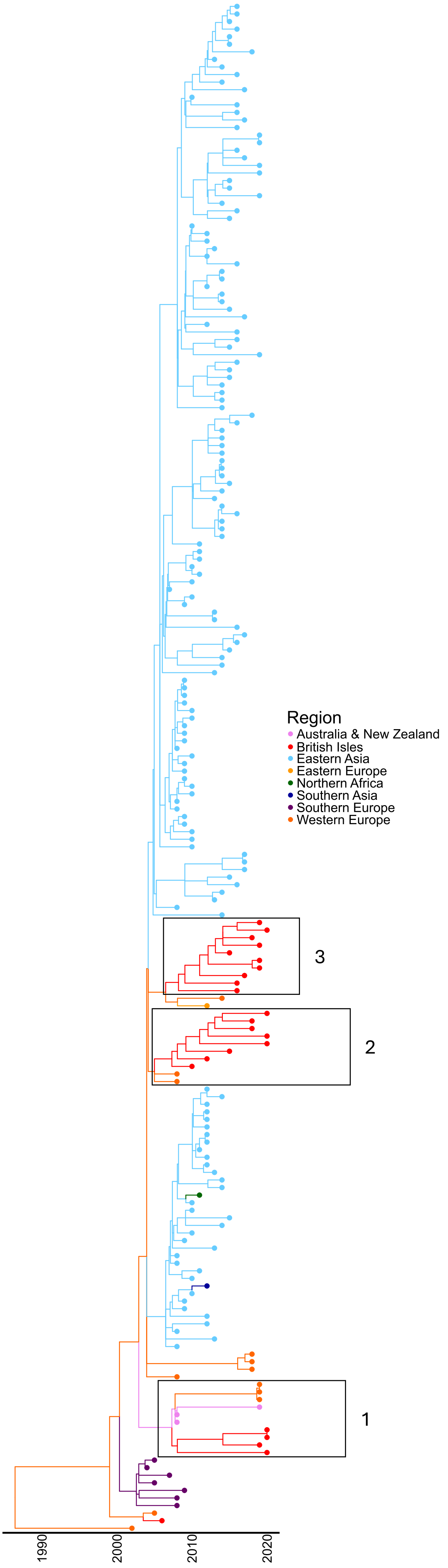


**S3 PPMV-1 genotype VI.2.1.1.2.2 incursion into tish Isles**. Treetime and mugration analysis of M-L eny of isolates identified as genotype VI.2.1.1.2.2. eny is determined by year of collection. Tips and es are coloured by region of origin.
