## Supplementary tables for "Phylogenetic analysis of pigeon paramyxovirus type 1 (PPMV-1) detected in the British Isles between 1983 – 2023": Supplementary tables - Byrne et al.pdf

**Supplementary Table 1: Coverage statistics from whole genome sequencing (WGS) of pigeon paramyxovirus-1.** Quality assessment of the WGS sequence obtained. For each sequence the total reads, genome coverage and mean depth of reads were obtained using samtools.

| Strain | Number reads | Coverage | Mean depth | Sequencing Method |
|---|---|---|---|---|
| APMV-1/Pigeon/United_Kingdom/AV1521/1983 | 485389 | 100 | 4214.52 | Illumina |
| APMV-1/Pigeon/United_Kingdom/AV1368/1984 | 2303364 | 100 | 19484.2 | Illumina |
| APMV-1/Pigeon/United_Kingdom/AV0868/1986 | 3364268 | 100 | 27963.2 | Illumina |
| APMV-1/Pigeon/United_Kingdom/AV0837/1988 | 4523873 | 100 | 38612.8 | Illumina |
| APMV-1/Pigeon/United_Kingdom/AV1528/1989 | 5873152 | 100 | 47429.5 | Illumina |
| APMV-1/Chicken/United_Kingdom/AV0710/1991 | 3562386 | 100 | 30737.2 | Illumina |
| APMV-1/Fantail/United_Kingdom/AV0250/1996 | 5629808 | 100 | 44403.5 | Illumina |
| APMV-1/Pigeon/United_Kingdom/AV0699/1997 | 4997754 | 100 | 39425.1 | Illumina |
| APMV-1/Pigeon/Ireland/AV0802/2000 | 4035709 | 100 | 34683.7 | Illumina |
| APMV-1/Pigeon/United_Kingdom/AV0771/2000 | 3186217 | 100 | 28005 | Illumina |
| APMV-1/Pigeon/Ireland/AV0931/2001 | 4518094 | 100 | 38368.9 | Illumina |
| APMV-1/Ireland/AV0974/2002 | 3418694 | 100 | 28167.2 | Illumina |
| APMV-1/Pigeon/United_Kingdom/AV0223/2002 | 4686169 | 100 | 39497.9 | Illumina |
| APMV-1/Pigeon/United_Kingdom/AV0956/2003 | 5408102 | 100 | 44120.9 | Illumina |
| APMV-1/Pigeon/United_Kingdom/AV3039/2004 | 3970965 | 100 | 33938.6 | Illumina |
| APMV-1/Pigeon/United_Kingdom/AV0222/2005 | 4706166 | 100 | 39556.2 | Illumina |
| APMV-1/Partridge/United_Kingdom/AV7575/2006 | | | | Sanger |
| APMV-1/Dove/United_Kingdom/AV7487/2006 | 5104102 | 100 | 42620 | Illumina |
| APMV-1/Pigeon/United_Kingdom/AV1875/2007 | 4485133 | 100 | 37969.6 | Illumina |
| APMV-1//United_Kingdom/AV2278/2007 | 5879077 | 99.9934 | 46804.4 | Illumina |
| APMV-1/Pigeon/Ireland/AV1521/2008 | 891481 | 100 | 8034.01 | Illumina |
| APMV-1/Pigeon/United_Kingdom/AV1271/2009 | 739706 | 100 | 6772.86 | Illumina |
| APMV-1/Pigeon/United_Kingdom/AV0257/2010 | 6075574 | 100 | 48194.1 | Illumina |

| | | | | |
|---|---|---|---|---|
| APMV-1/Pigeon/Ireland/AV1262/2011 | 163814 | 99.9605 | 1527.1 | Illumina |
| APMV-1/Pigeon/United_Kingdom/AV0104/2011 | 3376145 | 100 | 29338.5 | Illumina |
| APMV-1/Pigeon/United_Kingdom/AV0682/2013 | 4461728 | 100 | 38172.5 | Illumina |
| APMV-1/Pigeon/United_Kingdom/AV0593/2015 | 29107 | 98.3741 | 268.767 | Illumina |
| APMV-1/Pigeon/United_Kingdom/AV1419/2018 | 273133 | 100 | 2557 | Illumina |
| APMV-1/Dove/United_Kingdom/AV0229/2018 | 190703 | 100 | 1786.87 | Illumina |
| APMV-1/Pigeon/United_Kingdom/AV0151/2019 | 1348905 | 100 | 12341.7 | Illumina |
| APMV-1/Pigeon/United_Kingdom/AV0170/2019 | 713254 | 100 | 6513.69 | Illumina |
| APMV-1/Pigeon/United_Kingdom/AV1652/2020 | | | | Sanger |
| APMV-1/Pigeon/Guernsey/AV0310/2021 | 22559 | 96.8141 | 205.903 | Illumina |
| APMV-1/Pigeon/United_Kingdom/AV0167/2021 | 12607 | 97.9134 | 115.864 | Illumina |
| APMV-1/Pigeon/United_Kingdom/AV0489/2021 | | | | Sanger |
| APMV-1/Pigeon/United_Kingdom/AV01837/2021 | 29329 | 98.5321 | 273.07 | Illumina |
| APMV-1/Pigeon/United_Kingdom/AV01902/2021 | 2581655 | 98.4347 | 22687.8 | Illumina |
| APMV-1/Pigeon/United_Kingdom/AV04178/2022 | 33063 | 99.9603 | 208.167 | Illumina |
| APMV-1/Feral_Pigeon/United_Kingdom/AV0533/2022 | 28871 | 98.2425 | 265.419 | Illumina |
| APMV-1/Feral_Pigeon/United_Kingdom/AV0546/2022 | 28799 | 98.4071 | 267.709 | Illumina |
| APMV1/Pigeon/United_Kingdom/AV05158/2022 | 6874 | 99.3286 | 41.4537 | Illumina |
| APMV1/Collared_Dove/United_Kingdom/AV5664/2022 | 165650 | 100 | 155750 | ONT |
| APMV1/Rock_Dove/United_Kingdom/AV5833/2023 | | | | Sanger |
| APMV-1/Pigeon/United_Kingdom/AV05928/2022 | | | | Sanger |
| APMV1/Pigeon/United/Kingdom/53/2023 | 354591 | 99.8552 | 2233.93 | Illumina |
| APMV1/Rock_Dove/United_Kingdom/AV0101/2023 | | | | Sanger |
| APMV1/Dove/United_Kingdom/AV0153/2023 | | | | Sanger |
| APMV1/Starling/United_Kingdom/AV0676/2023 | 2338 | 100 | 1149.41 | ONT |
| APMV1/Rock_Dove/United_Kingdom/AV0677/2023 | 213889 | 100 | 192144 | ONT |

| | | | | |
|---|---|---|---|---|
| APMV1/Rock_Dove/United_Kingdom/AV1501/2023 | 131105 | 100 | 123927 | ONT |
| APMV-1/Chicken/Abu_Grab/Iraq/1968 | | | | Sanger |

**Supplementary Table 2: Log marginal likelihood of distinct clock and tree models for BEAST analysis.**

| | **P.S.** | **S.S.S.** |
|---|---|---|
| **Relax/Bayesian SkyRide** | -13135.7043530497 | -13142.4365241774 |
| **Relax/Constant Population** | -13141.9564712257 | -13148.1396438063 |
| **Strict/Bayesian SkyRide** | -13191.7540223194 | -13199.8483479336 |
| **Strict/Constant Population** | -13175.8157193644 | -13181.403039191 |

| | **P.S.** | **S.S.S.** |
|---|---|---|
| **Relax/Bayesian SkyRide/Asymmetric** | -13167.0554443881 | -13168.6608443862 |
| **Relax/Bayesian SkyRide/Symmetric** | -13160.691471337 | -13169.8571527129 |

**Supplementary Table 3: Bayes factor and posterior probabilities following SpreaD3 analysis of VI.2.1.1.2.2 transmissions.**

| FROM | TO | BAYES FACTOR | POSTERIOR PROBABILITY |
|---|---|---|---|
| Eastern Asia | Western Europe | 0.130649028 | 0.047161427 |
| Eastern Asia | Eastern Europe | 0.267238186 | 0.09193423 |
| Eastern Asia | Northern Africa | 0.233145498 | 0.081157649 |
| Eastern Asia | Southern Europe | 0.243953824 | 0.084601711 |
| Eastern Asia | Southern Asia | 0.250618573 | 0.086712587 |
| Eastern Asia | United Kingdom | 0.740545841 | 0.219086768 |
| Eastern Asia | Australia and New Zealand | 23.58453051 | 0.899344517 |
| Western Europe | Eastern Europe | 0.136961978 | 0.049327852 |
| Western Europe | Northern Africa | 182.255843 | 0.985723808 |
| Western Europe | Southern Europe | 131.9726622 | 0.980391068 |
| Western Europe | Southern Asia | 0.121153045 | 0.043884013 |
| Western Europe | United Kingdom | 0.126295314 | 0.045661593 |
| Western Europe | Australia and New Zealand | 223.6372046 | 0.988334629 |
| Eastern Europe | Northern Africa | 0.247808612 | 0.085823797 |
| Eastern Europe | Southern Europe | 0.243603899 | 0.084490612 |
| Eastern Europe | Southern Asia | 0.277588532 | 0.095156094 |
| Eastern Europe | United Kingdom | 0.350463246 | 0.117209199 |
| Eastern Europe | Australia and New Zealand | 42.10439867 | 0.941006555 |
| Northern Africa | Southern Europe | 0.3213973 | 0.108543495 |

| | | | |
|---|---|---|---|
| Northern Africa | Southern Asia | 0.237842374 | 0.082657483 |
| Northern Africa | United Kingdom | 0.213996143 | 0.074991668 |
| Northern Africa | Australia and New Zealand | 0.238888211 | 0.082990779 |
| Southern Europe | Southern Asia | 0.246931619 | 0.08554605 |
| Southern Europe | United Kingdom | 0.208181546 | 0.073102989 |
| Southern Europe | Australia and New Zealand | 0.238016627 | 0.082713032 |
| Southern Asia | United Kingdom | 0.227598648 | 0.079380069 |
| Southern Asia | Australia and New Zealand | 117.6594594 | 0.978057994 |
| United Kingdom | Australia and New Zealand | 47515.48996 | 1 |
